# Melatonin Protects Against Palmitate-Induced Lipid Accumulation and Leptin Resistance via PI3K/AKT in 3T3-L1 Cell line

**DOI:** 10.1101/2025.03.24.640716

**Authors:** Vennila Suriyagandhi, Ramkumar Arunachalam, Vasanthi Nachiappan

**Affiliations:** Biomembrane Lab, Department of Biochemistry, School of Life Sciences, Bharathidasan University, Tiruchirappalli-620 024, Tamilnadu, India; Department of Environmental Biotechnology, School of Environmental Sciences, Bharathidasan University, Tiruchirappalli-620 024, Tamilnadu, India

**Keywords:** Melatonin, Palmitate, Melatonin receptor (MT1 & MT2), Obesity, Lipid accumulation, Leptin resistance

## Abstract

**Background:** A high-fat diet (HFD) (which contains palmitate) is the primary factor for diet-induced obesity. Lack of sleep leads to the selection of HFD to maintain wakefulness. Sleep deprivation, along with HFD, induces obesity by leptin resistance. The sleep hormone melatonin secreted at night is responsible for the sleep-wake cycle based on circadian rhythm. It has various other functions in the central and peripheral axes, including food intake and energy expenditure. Primarily, melatonin regulates intracellular signaling via its receptors MT1 and MT2 (GPCR) in the cell membrane and can directly enter the cytoplasm.

**Aim:** To study the role of melatonin and its receptor-mediated signaling in palmitate-induced obesity and leptin resistance using the 3T3-L1 cell line.

**Methods:** Lipid accumulation was induced in differentiated 3T3-L1 cell lines by sodium palmitate (0.8 mM). Further, we exposed the cells to melatonin (1 mM) and melatonin receptor inhibitors (luzindole/and 4P-PDOT). Lipid accumulation and melatonin’s effect were studied using thin-layer chromatography and oil-red O staining, and the mRNA and protein expression of lipid-synthesizing genes and lipases were analyzed using qRT-PCR and western blotting. Leptin secretion was quantified by ELISA assay. Calcium imaging was done by Fura2-AM fluorescent staining.

**Results:** Palmitate induction in differentiated 3T3-L1 cell line elevated triacylglycerol (TAG) in lipid droplets along with an increase in the mRNA expression of PPARγ, SREBP1c, FASN, ACC1, and leptin secretion but decreased the lipases (HSL and ATGL) and PI3K and AKT. Treatment with melatonin restored the TAG level, lipid droplet size, and gene expression of PPARγ, SREBP1c, FASN, and ACC1, as well as leptin secretion, but increased protein expression of lipases. Melatonin receptor inhibitors, along with palmitate and melatonin, reversed the positive effects of melatonin on fat accumulation and leptin resistance. Intracellular calcium levels were reduced with palmitate induction, and melatonin rescued the calcium level equal to the control group through its receptor signaling. Melatonin restores the mRNA expression of PI3K and AKT genes in palmitate-induced conditions via its receptors.

**Conclusion:** Melatonin is vital for hormonal regulation and energy metabolism, including obesity and leptin secretion. Studying the role of melatonin in leptin regulation and lipid metabolism will help us combat the pathologies of obesity caused by HFD. From our study, the action of melatonin against lipid accumulation and the regulation of leptin secretion is mediated through its receptors MT1 and MT2 by activating the downstream PI3k/AKT signaling.

**Highlights:**

- Melatonin restores palmitate-induced lipid accumulation via its receptors.
- Melatonin reduces lipogenesis and increases lipolysis in lipid-accumulated adipocytes.
- Melatonin rescues palmitate-induced leptin resistance through its receptors by altering the intracellular calcium levels.
- Melatonin, via its receptors, restores palmitate-induced lipid accumulation by activating the PI3K/AKT signaling.

## 1. Introduction

Globally, obesity is the prime factor for developing non-communicable diseases like cardiovascular disease, type 2 diabetes, and some types of cancers. The spending for obesity treatment costs around $2 trillion annually in 2019. The primary reasons for obesity development are lack of physical activity, a high-fat diet, consumption of processed foods, and eating at the wrong time [1]. The nutritional overload in diet-induced obesity causes enormous expansion of adipose tissue, resulting in inflammation [2].

Adipose tissue is known as the body’s energy reservoir and stores energy as TAG in lipid droplet-LD. The LDs expand the cells, accommodate extra lipids, and are responsible for the differentiation of preadipocytes into mature adipocytes [3]. In addition, adipocytes secrete protein hormones that act as endocrine, paracrine, and autocrine. The adipocyte hormones leptin and adiponectin send signals to the brain regarding energy availability and regulate energy expenditure [4]. Leptin sends signals to the hypothalamus via its transporters, restricts food intake, and induces energy expenditure by activating JAK-STAT signaling through its receptor [5, 6].

With the current sedentary lifestyle, excess food intake and reduced energy expenditure result in fat accumulation in the adipocytes, which produces inflammatory cytokines [7].

In addition, accumulation of excess fat by diet-induced obesity increases leptin secretion and leptin transport to the hypothalamus. Uncontrolled secretion of excess leptin saturates the leptin transporter and its receptor activation, unable to regulate energy expenditure and food intake, and results in leptin resistance [8, 9, 10]. Palmitate is the primary fatty acid in a high-fat diet (HFD), which enhances obesity and other related diseases. In addition, HFD induces dysfunction of adipocytes, causes inflammation and insulin resistance, and dysregulates energy balance in mice and humans [11]. In addition, excess palmitate in the circulation is reported to lead to autophagy and ER stress.

Research shows that altered circadian rhythms caused by sleep deprivation increase food intake, decrease energy expenditure, and result in obesity [12]. Another study reported that sleep restriction leads a person to select high-calorie foods along with alteration in appetite-regulating hormones like leptin, ghrelin, and PYY compared to their secretion during normal circadian rhythm [13]. The strategy to reduce obesity is to use anti-obesity drugs. Effective anti-obesity drugs are available to reduce de novo lipogenesis, increase lipolysis, and suppress adipose tissue expansion [14]. The use of commercially available anti-obesity medications for a longer time and higher dosage can lead to unacceptable side effects [15]. Hence, we selected melatonin as an anti-obesity agent, which has fewer side effects compared to commercially available drugs. Melatonin is a pineal gland sleep hormone secreted in the dark, sending its signal all over the body to maintain the day and night cycle [16]. Besides serving as the sleep hormone, melatonin also possesses antioxidant properties, scavenges free radicals, and has anti-inflammatory and anti-apoptosis properties [17]. Our previous in silico analysis predicted that melatonin targets various inflammatory cytokines and AKT1 to control leptin resistance-induced obesity [18].

Melatonin manifests its cellular impact by coupling to its high-affinity receptors MT1 and MT2. These receptors have seven transmembrane-spanning G-proteins and are present in the plasma membrane. In addition, melatonin diffuses across the morpho-physiological barriers without its receptors and reaches the intracellular region, including the blood-brain barrier and placenta [19]. The G protein acts as a molecular switch in the G protein-coupled receptors and modulates intracellular signaling. Upon activation, the MT1 receptor inhibits forskolin-stimulated cAMP, PKA signaling and phosphorylation of CREB and activation of PI3k signaling, mitogen-activated protein kinase 1/2, extracellular signal-regulated kinase 1/2, and potassium channels by coupling with both pertussis toxin-sensitive Gi and pertussis toxin insensitive Gq/11 G proteins. Activation of MT2 receptor inhibits cAMP, cGMP and activates protein kinase C (PKC) and PLA2 [20]. Further, these effectors regulate downstream PKA, PLC, IP3, PI3k, and AKT [21, 22]. The activated IP3, PLC, and PIP2 release calcium from the endoplasmic reticulum and are responsible for various cellular processes, particularly cell survival and cell death [23]. Our previous review explained that melatonin treatment reduces adipocyte leptin secretion in high-fat-fed/obese animals [24].

However, the exact mechanism of melatonin signaling in the obese models and its role in lipid accumulation and leptin secretion is still unclear. Therefore, we wanted to understand the role of melatonin and the involvement of melatonin signaling via its receptors MT1 and MT2 in obesity. In this study, we examined the role of melatonin signaling lipid accumulated state to analyze the lipid metabolism and the leptin level in the 3T3-L1 cell line using MT1 and MT2 selective antagonists luzindole and 4P-PDOT. We found that melatonin has lipid-lowering and leptin regulatory effects and was achieved via the MT1 and MT2 receptor signaling in obese cell models.

## 2. Materials and Methods

### 2.1. Cell culture and differentiation

The 3T3-L1 fibroblast was obtained from NCCS, Pune, India. This preadipocyte is maintained in Dulbecco’s Modified Eagle medium (DMEM, 4.5 g/L glucose, with 10% Fetal bovine serum (FBS) (Gibco™-A3160502), 1% Antibiotic-Antimycotic (Gibco™-15240096) in a humidified incubator with 5% CO2 until full confluence. The cell line is then supplemented with a differentiation medium (1 mM insulin, 500 µM 3-isobutyl-1-methylxanthine, 500 nM dexamethasone) for two days. Later, the medium was switched to a maintenance medium containing DMEM, FBS, and 1 mM insulin (I9278, Sigma Aldrich, India) with alternate days of medium change for up to 15 days until the adipocytes were differentiated to mature ones. Maturation was confirmed by visual inspection and Oil Red O staining (data not included).

### 2.2. Cell viability by MTT Assay

Briefly, after differentiation and serum starvation, the cells in a 96-well plate were treated with various melatonin concentrations for 12 h. The medium was removed, and to each well, 10 μl of MTT solution (5 mg/ml) (M5655, Sigma Aldrich, India) was added and incubated for 3 h. Later, the MTT was removed, and DMSO was added to solubilize the crystals. The absorbance was measured using a microplate reader (Bio-Rad) set at 450 nm wavelength.

### 2.3. Sodium palmitate, melatonin, and melatonin inhibitors treatment

The differentiated adipocyte cells were washed with PBS (pH 7.4), serum starved for 24 h, and induced with 5% fatty acid-free BSA conjugated 0.8 mM sodium palmitate for 12 h. 1 mM Melatonin (1380105, Sigma Aldrich, India) (dissolved in 0.25% DMSO) along with melatonin receptor inhibitors 10 µM 4P-PDOT (SML1189, Sigma Aldrich, India) and/or 10 µM luzindole (L2407, Sigma Aldrich, India) added to the cells were incubated for another 12 h. This treatment is used for the following analysis.

### 2.4. Oil Red O staining for lipid accumulation analysis

After the treatment mentioned above, the cells were fixed with 4% paraformaldehyde, washed with 1X PBS, and stained with Oil Red O (O1391. Sigma Aldrich, India) (0.5% freshly prepared) staining solution for 3 h. Again, the cells were washed with 1X PBS, and microscopic pictures were taken using an inverted microscope with 20 X magnification. The bound oil red o stain in the cells of each well was eluted with 100 μl isopropanol and transferred to a new 96-well plate. Absorbance was read at 490 nm in a microplate reader (Bio-Rad) to measure total lipid content indicative of neutral lipids, including TAG.

### 2.5. Lipid extraction and ‘Thin Layer Chromatography – TLC’

Briefly, after treatment, as mentioned above, cells were collected and centrifuged. After centrifugation, the culture medium was removed, and cells were solubilized with RIPA lysis buffer. Protein was quantified by Lowry’s method [25], and samples (cell lysates) with equal protein concentrations were used for lipid analysis. Lipid extraction was done following the Bligh and Dyer method [26] with slight modifications. Briefly, to the cell lysate, a 2:1 ratio of chloroform and methanol was added and vortexed. Then, an equal volume of acidified water (2 % ortho-phosphoric acid) was added and vortexed vigorously. The neutral lipids were separated by TLC using petroleum ether, diethyl ether, and acetic acid (17.5:7.5:0.200, v/v) as the solvent system. Individual lipids were identified by comparing the standard’s Rf values with the unknown’s Rf values. The TLC spots were exposed to iodine in a glass chamber, and once the lipids were sufficiently visible, the plates were removed and quantified using a densitometry scanner.

### 2.6. Leptin ELISA Assay

After the above-mentioned treatments, the cell culture medium was collected and stored at -80 °C for further use. The secreted leptin level was measured by a mouse Leptin ELISA kit (RAB0334. Sigma Aldrich, India) according to the manufacturer’s protocol using the sandwich ELISA method. The absorbance at 450 nm was measured in a Bio-Rad microplate reader.

### 2.7. RNA isolation and gene expression analysis by qRT-PCR

RNA was isolated from the cells with the mentioned treatment using TRIZOL (T9424. Sigma Aldrich, India) and chloroform according to the manufacturer’s protocol. cDNA was constructed using two µg RNA with SuperScript™ IV First-Strand Synthesis System (Invitrogen™: 18091050). qRT-PCR was performed using SYBR Green (TB Green® Premix Ex Taq™ II (Tli RNase H Plus) TAKARA-RR820A) in Applied biosystems StepOne™ Plus Real-Time PCR system. The primers were designed using Primer-Blast (NCBI-NIH) (Table. 1), and the relative gene expression of *SREBP1c, ACC1, FASN, PPARγ, PI3K*, and *AKT* were quantified using the 2-ΔΔCt method [27], normalized with β-Actin that served as control.

Table 1: Mouse primer list used for qRT-PCR analysis. β-Actin gene used as control.

**Table 1:**
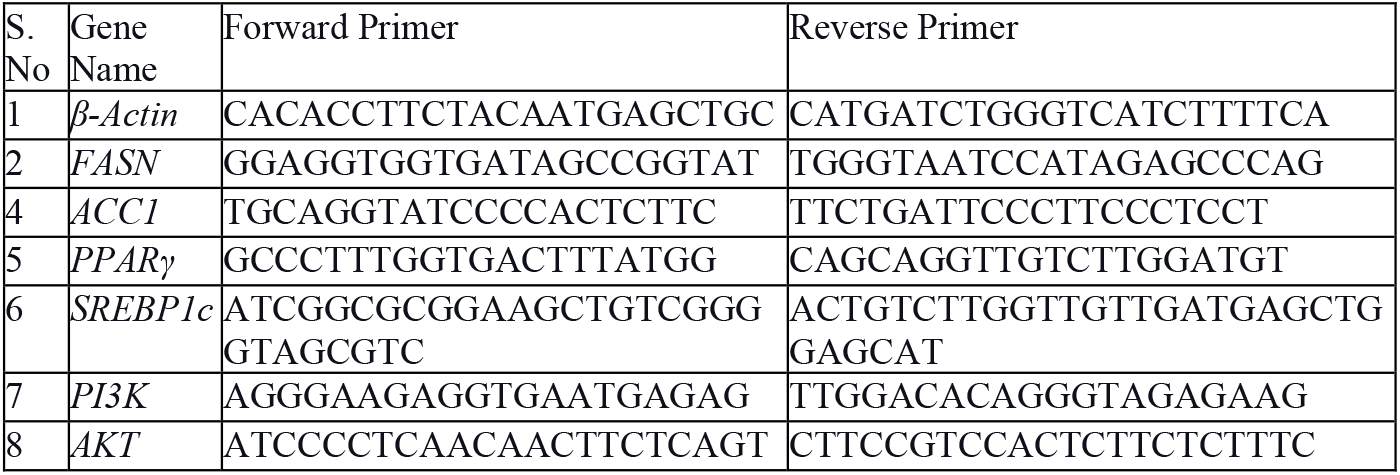
qRT-PCR mouse primer list.

### 2.8. Protein expression by Western Blotting

Cells after treatment were lysed with RIPA buffer, the lysed cells were centrifuged, and the supernatant was collected. Total protein analysis was done using Lowry’s method [25] with BSA as a standard. An equal concentration (40 µg) of protein was separated on 10-12% SDS PAGE and transferred to a nitrocellulose membrane. Furthermore, the membrane was treated with 5% blocking buffer (5% skimmed milk powder with 1X PBST) for 1.5 h at room temperature and incubated with a 1:1000 dilution primary antibodies (Cell signaling technology (HSL Antibody #4107, ATGL Antibody #2138, and β-Actin (8H10D10) Mouse mAb #3700)) overnight at 4 °C. Antibodies were removed, and the membrane was washed with 1X PBS and PBST three times for 5 min each. The membrane was exposed to ALP conjugated secondary antibody [(AP132A-Sigma-Aldrich, Goat Anti-Rabbit IgG Antibody, Alkaline Phosphatase conjugate, A3562-Sigma-Aldrich-Anti-Mouse IgG Alkaline Phosphatase antibody produced in goat)] with 2.5:5000 dilution for 1.5 h. The membrane was then washed with 1X PBS and PBST, and the bands were developed using the BCIP®/NBT Liquid Substrate System (Sigma Aldrich, India -B1911). The picture of the blots was taken with a digital camera. Bands on the blots were quantified using ImageJ software.

### 2.9. Calcium imaging using Fura-2 AM fluorescent staining

Briefly, the 3T3-L1 cells were seeded in a 24-well plate and differentiated for 15 days, and followed the above treatment, the existing medium was removed, the cells were washed with 1X PBS, and then Fura-2 AM (F0888-Sigma Aldrich, India) working solution (2 µM) was added to the wells, incubated for 20 min, and washed with 1X PBS. Immediately after washing, images were taken under a Carl Zeiss inverted fluorescent microscope with 20X magnification (excitation/emission at 340/510 nm).

### 2.10. Statistical Analysis

All data were expressed as mean ± S.E.M (standard error of the mean) from at least three different experiments. Data were analyzed using one-way ANOVA followed by Tukey’s multiple comparison test (GraphPad version 8.0). Significance was determined as a value of p<0.05 (*) and p<0.01 (**), p<0.001(***) and n.s. denotes non-significant.

## 3. Results

### 3.1. Cell viability by melatonin treatment

The viability of the differentiated 3T3-L1 cell line by melatonin was studied using an MTT assay. The differentiated 3T3-L1 cell line in 96 well plates was serum-starved for 24 h and then treated with various melatonin concentrations. Briefly, no significant growth defect was observed up to 1 mM melatonin concentration. Hence, 1 mM melatonin was used for further experiments (Fig. 1).

**Figure 1:**
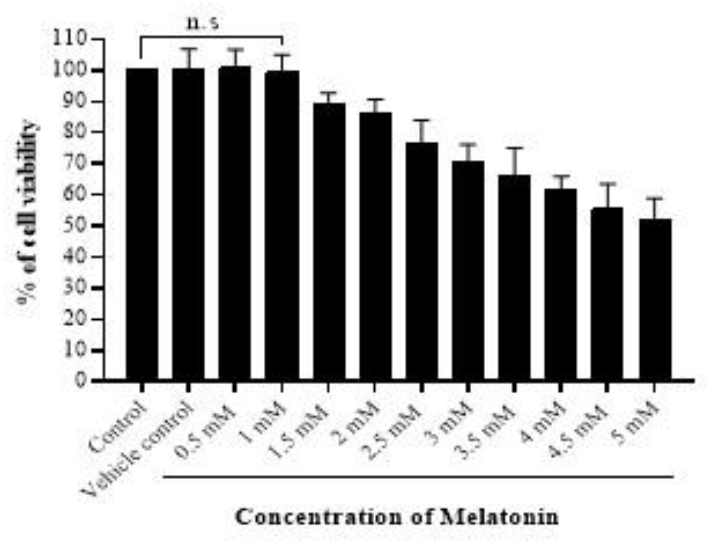
Cell viability of melatonin by MTT assay. Effect of melatonin on cell viability by different melatonin concentrations on the differentiated 3T3-L1 cell line. Melatonin from 0.5 mM to 5 mM added to the cells in a 96 well plate and incubated for 12 h. Formation of formazan crystals dissolved in DMSO were measured in a multiwell plate reader at 450 nm. Data is represented as % of viability vs control 100%. The statistical data is expressed as mean ± SE with significance *p<0.05, **p<0.01, ***p<0.001, n.s., not significant; n=3 represents three repeated independent experiments.

### 3.2. Melatonin dampens the lipid accumulation induced by palmitate

The lipid content of different treatment groups was analyzed by thin-layer chromatography. Based on previous literature [28], we used a 0.8 mM concentration of sodium palmitate (conjugated to 5% BSA) to induce obesity in the 3T3-L1 cell line. As expected, TAG was accumulated with palmitate induction. Further, the accrued TAG by palmitate was reversed by treating 1 mM melatonin. No inhibitors were added for this experiment. (Fig. 2).

**Figure 2:**
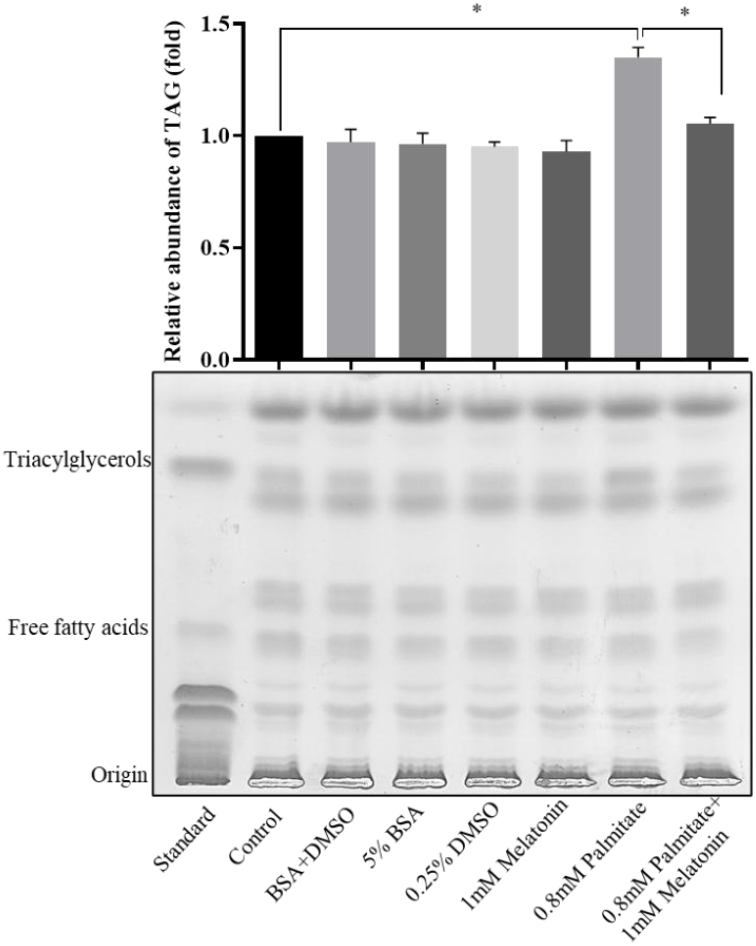
Analysis of TAG level with palmitate and melatonin treatment, along with vehicle controls treated to the differentiated 3T3-L1 cell line, was performed by thin-layer chromatography. To the differentiated cells, palmitate (0.8 mM conjugated to 5% BSA) was added and incubated for 12 h, melatonin (1 mM) was added and exposed for another 12 h. 5% BSA, and 0.25% DMSO was used as vehicle control with respective incubation—lipids extracted from the cell lysate by an equal amount of protein. Extracted lipid samples were loaded onto TLC plates. Individual lipids were identified by comparing the standard’s Rf values with the unknown’s Rf values. The bottom of the figure contains a picture of the TLC plate, and the top has a respective bar diagram of the TAG level. Data in the graph is represented as the relative abundance of TAG. The statistical data is expressed in mean ± SE. n=5 represents five repeated independent experiments.

### 3.3. Melatonin and its receptor signaling reduces the palmitate-induced intracellular lipid accumulation

Further, the palmitate-induced cells treated with melatonin were again exposed to melatonin receptor inhibitors (luzindole and 4P-PDOT) to study the role of melatonin and its receptors on lipid accumulation. The palmitate-induced cells depicted an increase of lipids in the lipid droplets (Fig. 3A), the lipid content including TAG compared to the control cells (Fig. 3B). In the palmitate-induced cells, melatonin treatment restored the accumulated lipid content in the lipid droplets to the control levels. On the contrary, melatonin failed to restore the accumulated lipid in the presence of melatonin receptor inhibitors, suggesting that melatonin acts via its receptor (MT1 and MT2) to reduce the intracellular lipids in the lipid droplets.

**Figure 3:**
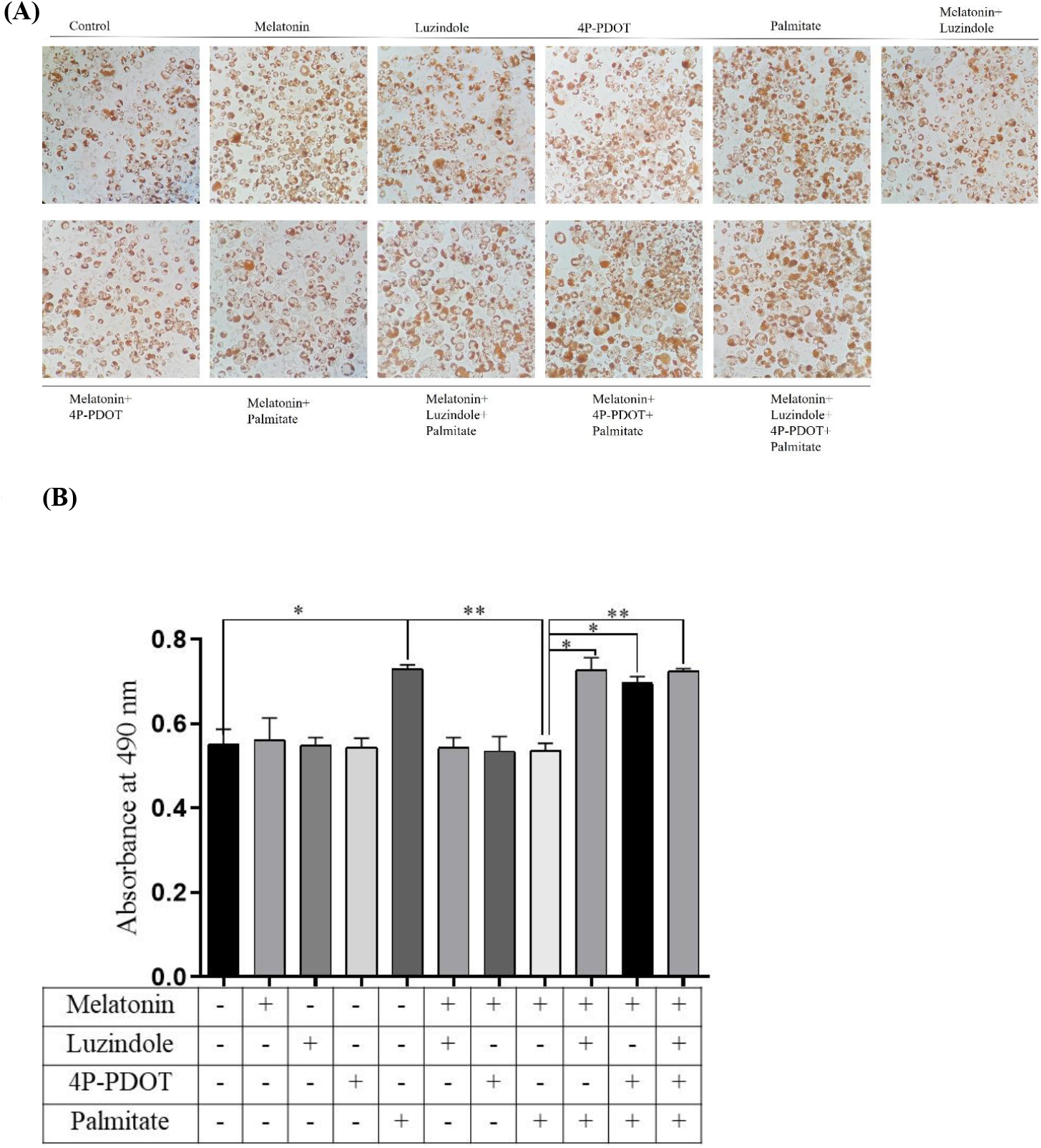
Oil Red O staining analysis of lipid droplets. Differentiated 3T3-L1 cells were exposed to various treatments. Lipid droplets were visualized by Oil Red O staining. Representative brightfield microscopic images (A) with 20X magnification were used to detect the lipid droplet/lipid content. Bar diagram (B) illustrates the absorbance of total lipid content indicative of neutral lipids including TAG by eluting the oil red o stain with isopropanol, measured in a multiwell plate reader at 490 nm absorbance. The statistical data is expressed in mean ± SE; with significance *p<0.05, **p<0.01, ***p<0.001, n.s., not significant; n=3 represents three repeated independent experiments.

### 3.4. Melatonin reduces the accumulated lipids by controlling lipogenesis and adipogenesis

Lipogenesis is the process of storing fatty acids in the form of TAG. Briefly, after differentiation, lipogenesis by palmitate and treatments (with melatonin/ receptor inhibitors) were determined using the mRNA expression of the lipid synthesizing genes *SREBP1c* (Fig. 4A), *ACC1* (Fig. 4B) and *FASN* (Fig. 4C) by qRT-PCR. The mRNA levels of lipogenesis genes were elevated in the palmitate-induced group. The melatonin treatment significantly reduced the *FASN, ACC1*, and *SREBP1c* mRNA expression. However, the melatonin receptor inhibitors elevated the gene expression of fatty acid synthesizing genes. The *PPARγ* gene is known as a master regulator of adipogenesis, and its expression was increased with palmitate induction. Melatonin significantly reduced the mRNA expression of *PPARγ* (Fig. 5) that was induced with palmitate, and the melatonin receptor inhibitors prevented the melatonin inhibition by increasing the *PPARγ* mRNA expression.

**Figure 4:**
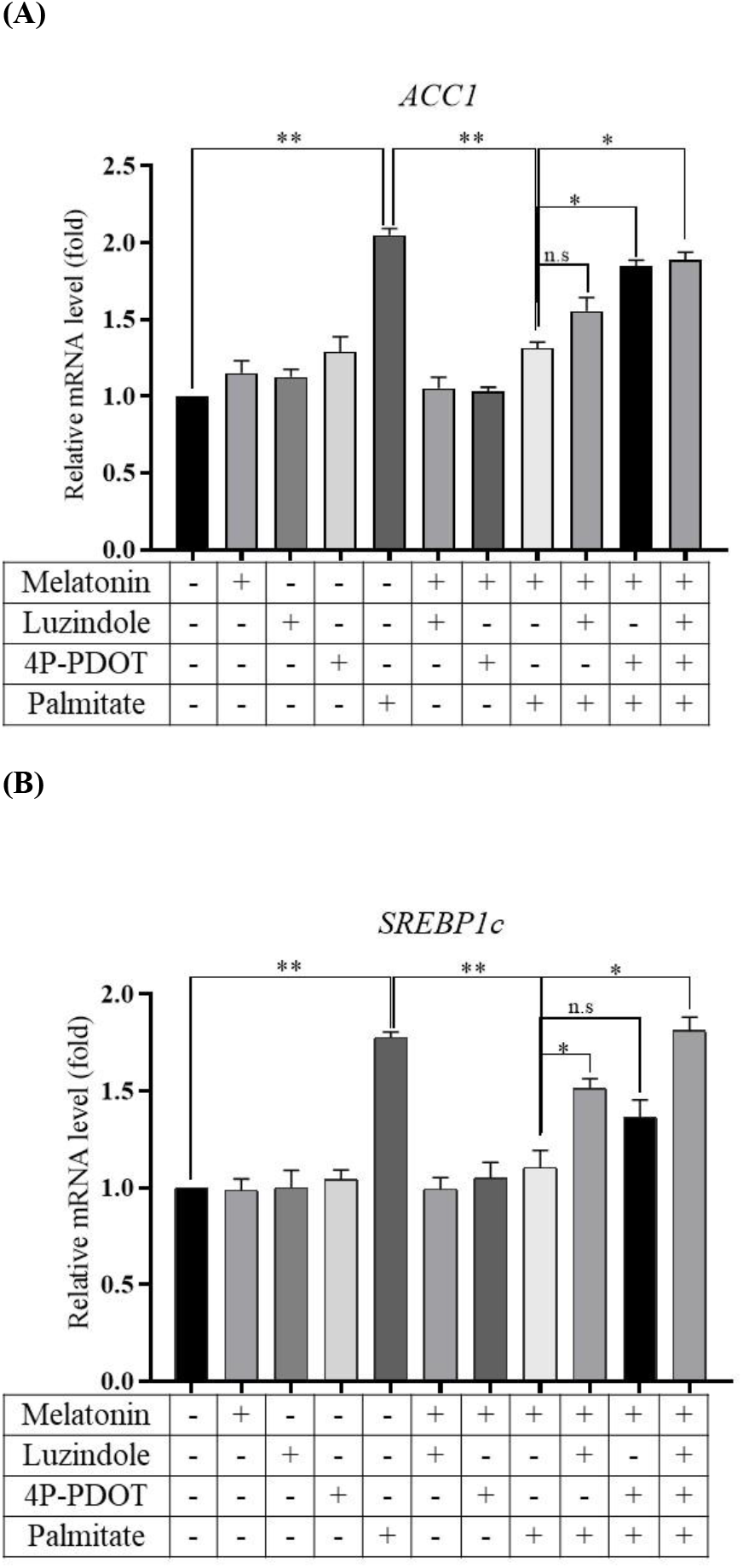
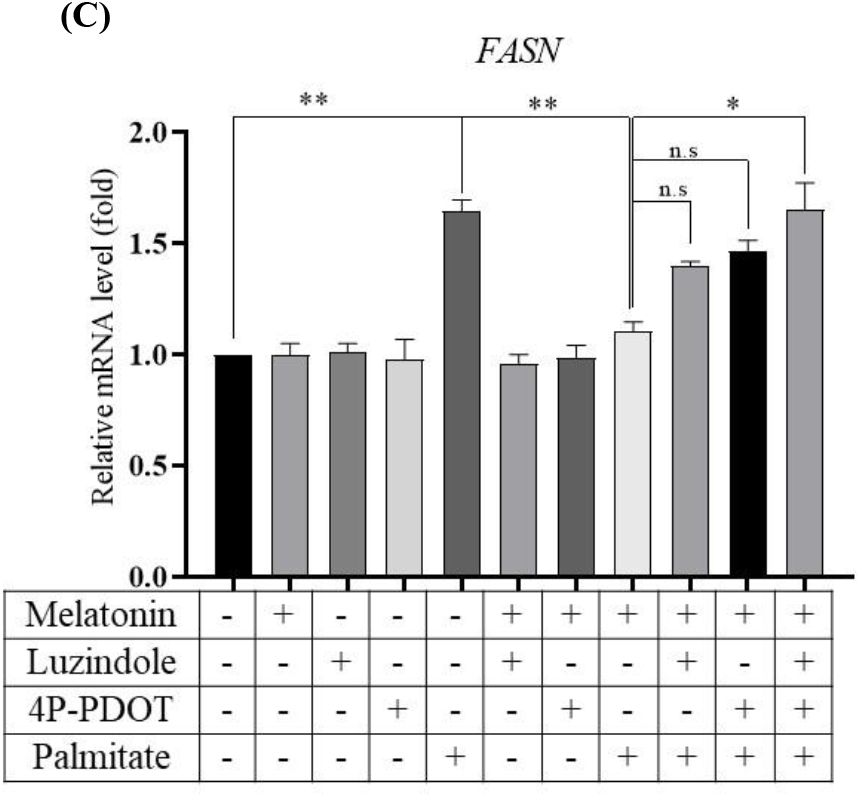
mRNA expression of genes involved in lipogenesis: The mRNA expression of lipogenesis genes was quantified using qRT-PCR. Differentiated 3T3-L1 cells were treated with palmitate (0.8 mM) for 12 h, melatonin (1 mM), and melatonin receptor inhibitors for an additional 12 h. The mRNA expression of (A) *SREBP1c*, (B) *ACC1*, and (C) *FASN* was quantified after the mentioned treatments. β-Actin was used as endogenous control. The statistical data is expressed in mean ± SE, with significance *p<0.05, **p<0.01, ***p<0.001, n.s., not significant; n=3 represents three repeated independent experiments.

**Figure 5:**
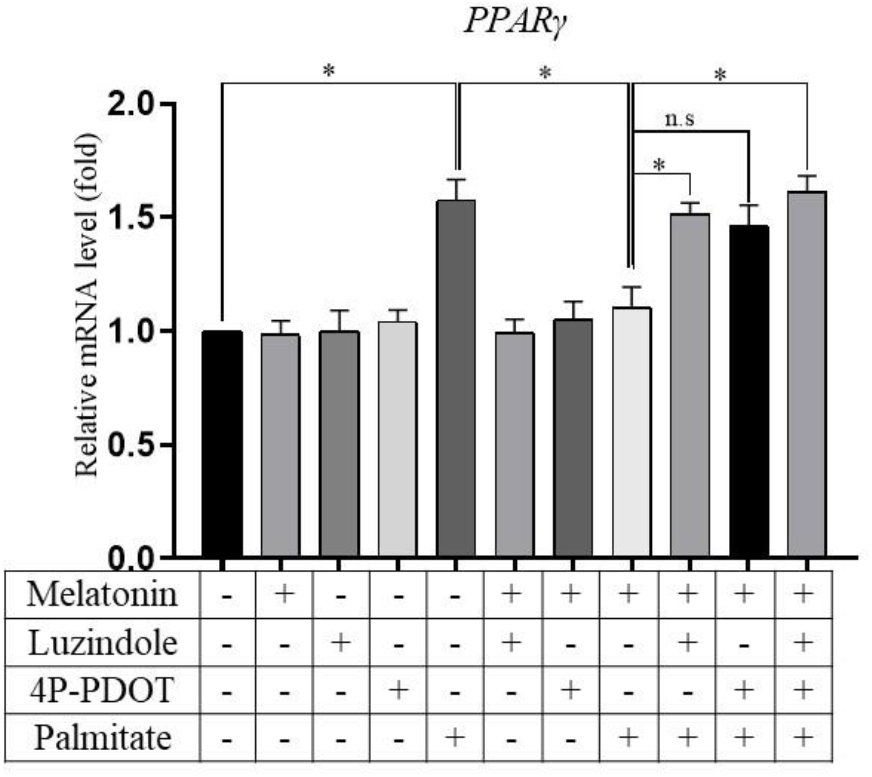
Effect of melatonin and its receptor inhibitors on the mRNA expression of adipogenesis gene PPARγ. The mRNA expression of *PPARγ* was quantified using qRT-PCR. Differentiated 3T3-L1 cells were exposed to palmitate, melatonin, and its receptor inhibitors. Expression of *PPARγ* was normalized by β-actin. The statistical data is expressed in mean ± SE; with significance *p<0.05, **p<0.01, ***p<0.001, n.s., not significant; n=3 represents three repeated independent experiments.

### 3.5. Melatonin regulates lipolysis via its receptor signaling

Lipolysis is essential for the maintenance of intracellular lipid levels, and during lipolysis, TAG is broken down into fatty acids and glycerol. The lipases ATGL and HSL generate energy from the storage lipids and are responsible for the TAG breakdown. The protein levels of lipase were reduced with palmitate induction compared to the control group, and melatonin rescued the reduced HSL and ATGL. However, the positive effect of melatonin on the lipases ATGL (Fig. 6A) and HSL (Fig. 6B) was blocked by melatonin receptor inhibitors, and we observed a reduction in the HSL and ATGL levels compared to the control and melatonin-treated groups (Fig. 6C).

**Figure 6:**
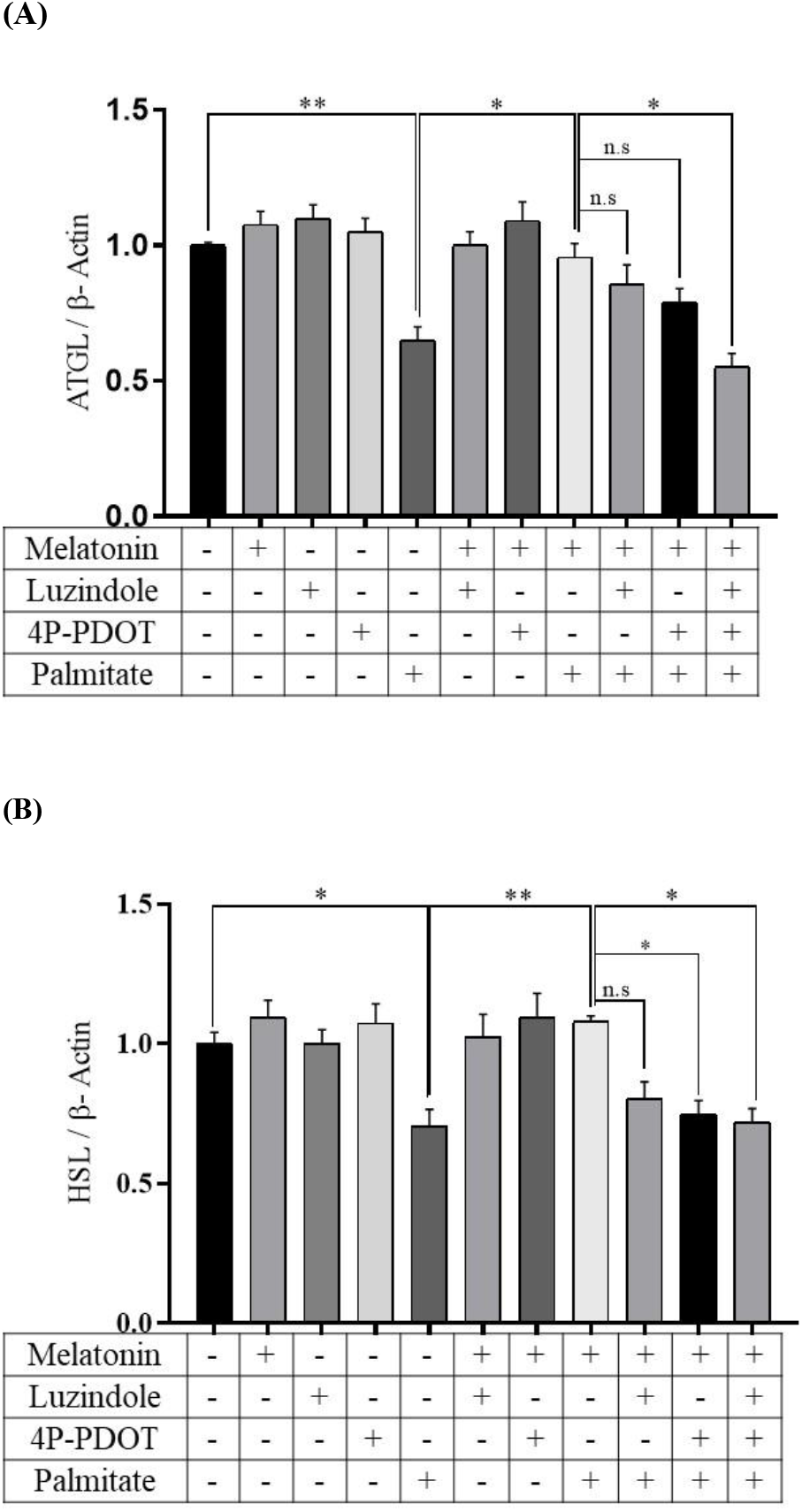
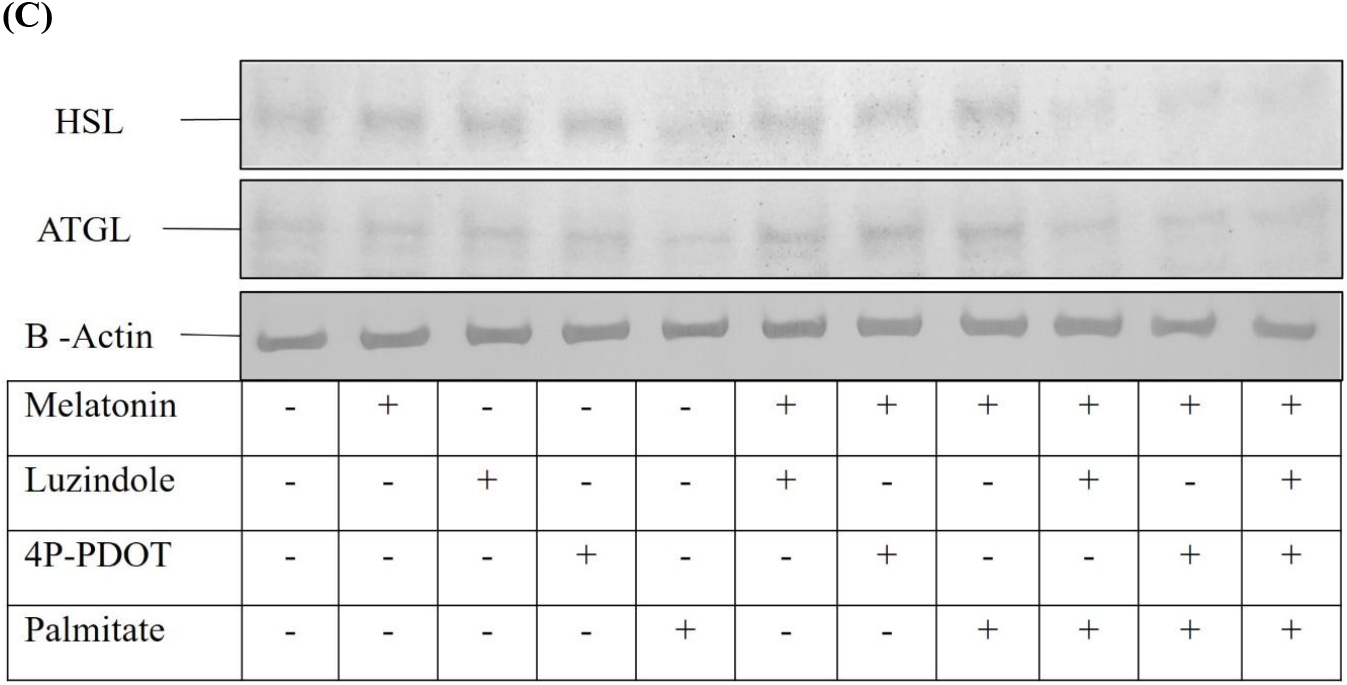
Effect of palmitate, melatonin, and its receptors on lipolytic enzymes: Representative bar diagram illustrates quantification of protein expression by Western blot analysis of (A) ATGL & (B) HSL on differentiated palmitate-induced 3T3-L1 cell line with melatonin and its receptor inhibitors. (C) Nitrocellulose membrane containing protein bands of HSL, ATGL, and β-Actin. The statistical data is expressed in mean ± SE, with significance *p<0.05, **p<0.01, ***p<0.001, n.s., not significant; n=3 represents three repeated independent experiments.

### 3.6. Melatonin rescues the cells from palmitate-induced leptin resistance by its receptor signaling

We used an ELISA assay to measure the secreted leptin level from the treated cells. Compared to the control group, palmitate induction significantly increased the leptin secretion in the culture medium, and melatonin treatment reduced the leptin level. Upon exposure to melatonin receptor inhibitors (single or both), the leptin secretion was increased as with the palmitate-treated group, which implies leptin secretion is based on melatonin via its receptors MT1 and MT2 (Fig. 7).

**Figure 7:**
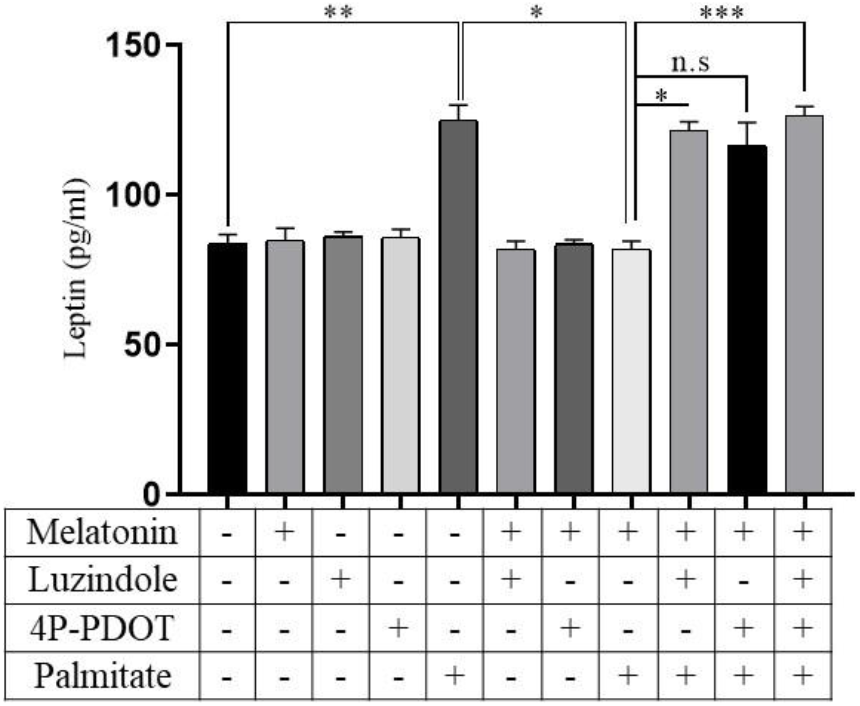
Role of melatonin receptor signaling in palmitate-induced leptin secretion. ELISA assay was used to measure the secreted leptin in cell culture medium from each experimental group with a sensitivity of 4 to 1000 pg/ml. The bar diagram depicts the pg/ml of leptin in cell culture medium with mentioned treatments. The concentration of leptin in the samples was compared against that of standard leptin. The statistical data is expressed in mean ± SE, with significance *p<0.05, **p<0.01, ***p<0.001, n.s., not significant; n=3 represents three repeated independent experiments.

### 3.7. Melatonin regulates calcium release to maintain leptin secretion by its receptor signaling

Intracellular calcium was visualized using Fura 2-AM fluorescent stain. The intracellular calcium level was significantly reduced with palmitate induction. However, palmitate with melatonin exposure restored the calcium level as in the control group. The palmitate-induced cells exposed to melatonin and its receptor inhibitors depicted a significant reduction in calcium levels similar to the palmitate induction alone (Fig. 8).

**Figure 8:**
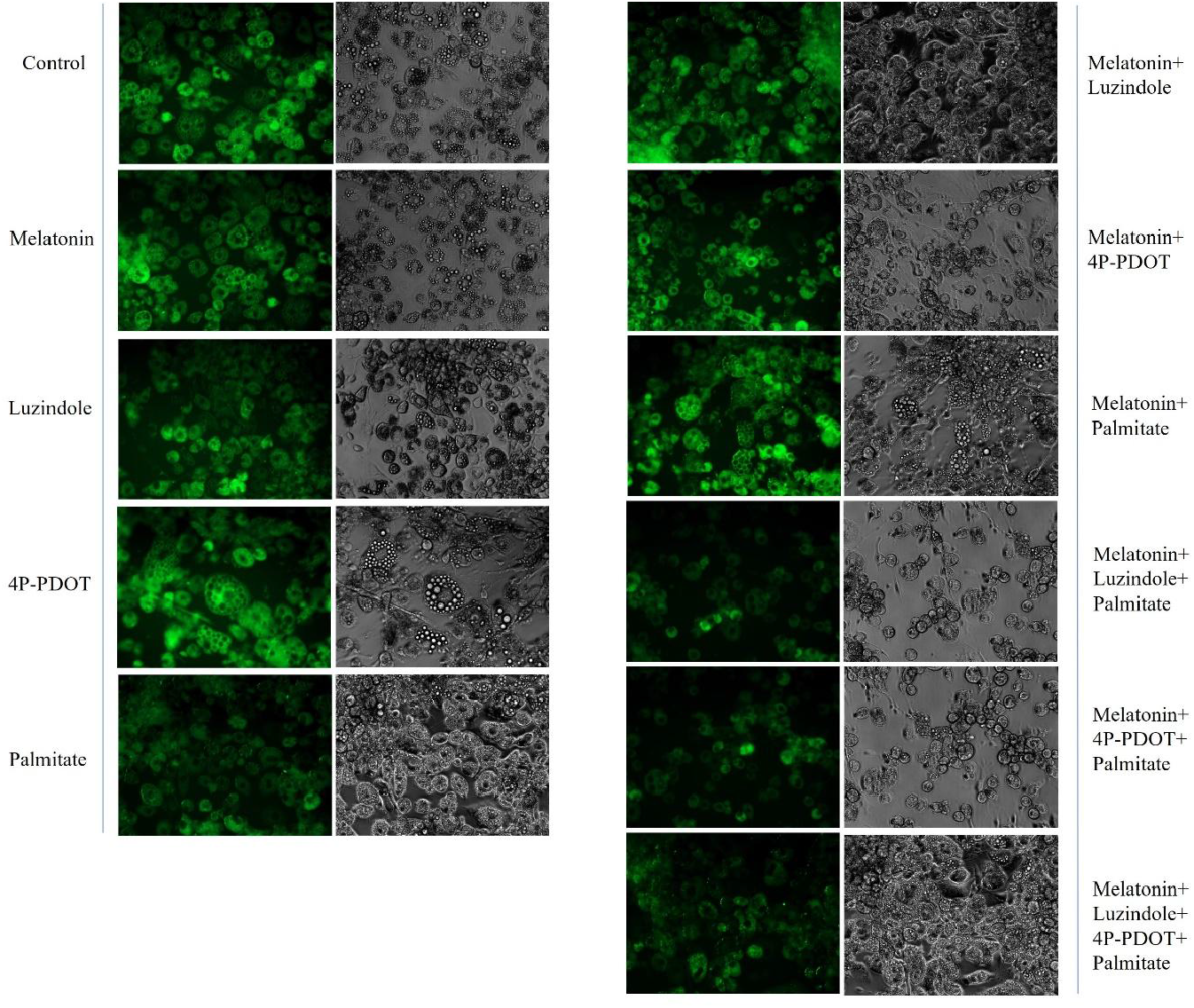
Effect of melatonin receptor signaling on intracellular calcium content. Fluorescent microscopy images showing the fluorescent intensity of Fura-2AM staining. Intracellular calcium content visualized on the differentiated palmitate-induced 3T3-L1 cell line with melatonin and its receptor inhibitors. Each treatment contains brightfield (right) and fluorescent (left) images. Microscopic images were taken with 20X magnification with a fluorescent inverted microscope; positions for each sample n=5 represent five repeated independent experiments.

### 3.8. The positive effect of melatonin in regulating lipogenesis and lipolysis via PI3k/AKT signaling

The signaling pathway that is regulated during lipid accumulation (due to palmitate induction) was analyzed by quantifying the mRNA expression of the *PI3K* and *AKT* genes. The gene expression of both *PI3K* (Fig. 9A) and *AKT* (Fig. 9B) was reduced in palmitate-induced cells, and melatonin exposure increased the mRNA expression of these genes, as observed with the control group level. However, the melatonin receptor inhibitors decreased the mRNA expression of *PI3K* and *AKT* genes and blocked the positive effect of melatonin signaling on lipid processing and leptin secretion by the PI3K/AKT pathway.

**Figure 9:**
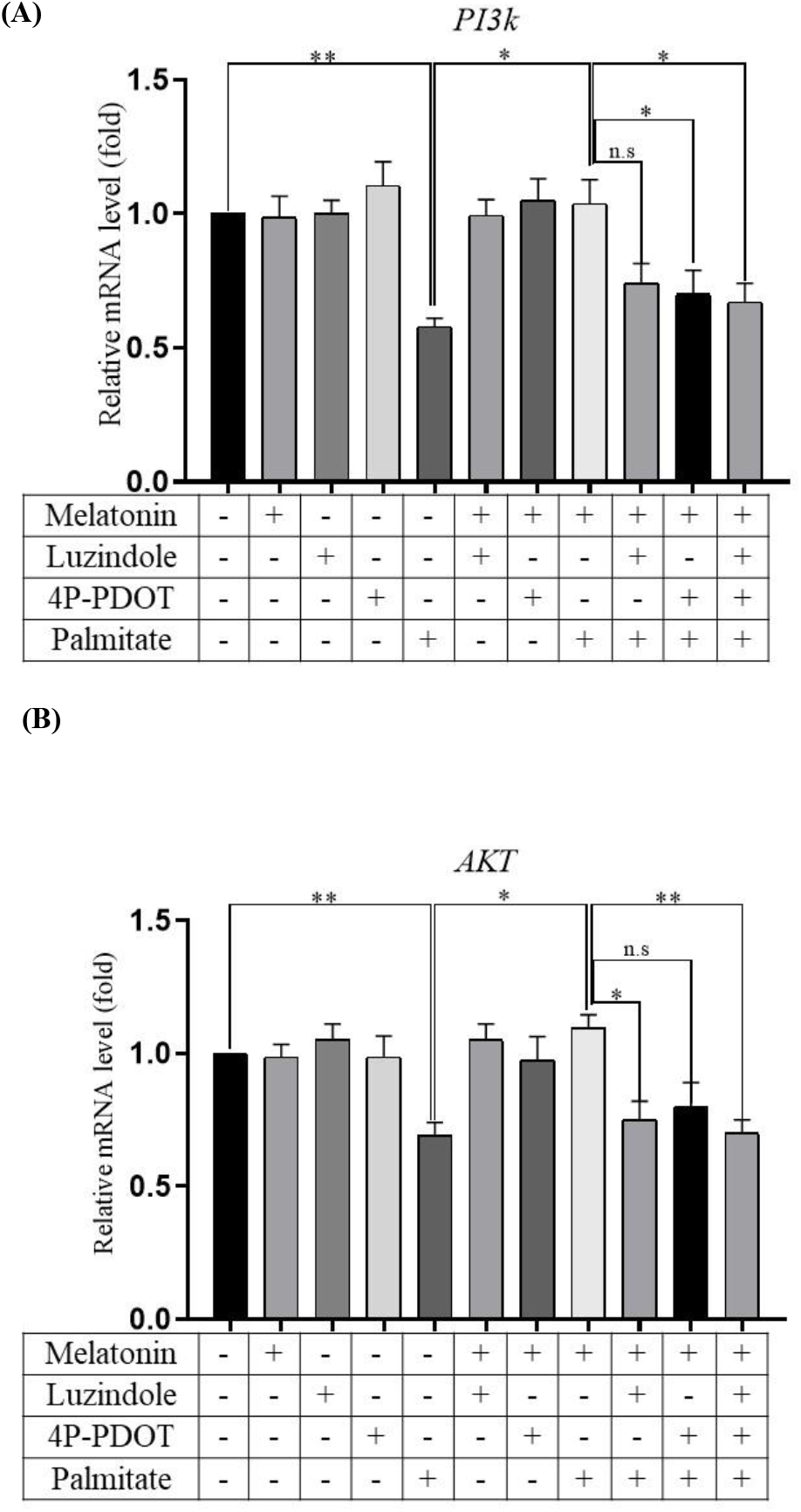
Effect of palmitate induction and role of melatonin and its receptors in lipid metabolism and leptin secretion by PI3K/AKT signaling. qRT-PCR was used to quantify the mRNA expression of the genes (A) PI3k and (B) AKT on differentiated palmitate-induced 3T3-L1 cell lines with melatonin and its receptor inhibitors. B-actin was used as an endogenous control. The statistical data is expressed in mean ± SE, with significance *p<0.05, **p<0.01, ***p<0.001, n.s., not significant; n=3 represents three repeated independent experiments.

## 4. Discussion

The mouse 3T3-L1 cell line is a widely used Invitro model system to study adipogenesis and lipid metabolism [29]. Dusko Lainscek et al. [30] suggested that an obese cell model is achieved with palmitate treatment in the 3T3-L1 cell line. Elevated levels of palmitate esters are observed in obese individuals. A previous report suggests that the obese model was created with the differentiated 3T3-L1 cells by treatment with sodium palmitate [31]. In this study, we used the 3T3-L1 cell line to study the role of melatonin and its receptor signaling in palmitate-induced lipid accumulation and leptin secretion.

As expected, the palmitate-induced cells accumulated lipids, particularly TAG, and were observed in the lipid droplets. Fig. 2 and 3 showed elevated neutral lipids, including TAG levels in lipid droplets with palmitate induction. It is reported that melatonin reduces obesity by decreasing inflammation and adipocyte hypertrophy in high-fat diet (HFD) fed mice [32]. Studies suggested that melatonin decreases serum TAG accumulation in HFD-induced hamsters and obese mice [33, 34, 35]. However, it is unclear how melatonin reduces obesity and increases energy expenditure. Therefore, we used melatonin and melatonin receptor inhibitors to investigate the role of melatonin signaling in leptin resistance and obesity management.

Briefly, the actions of melatonin by MT1 and MT2 receptors were blocked by the nonselective antagonist luzindole, which is competitive with melatonin. 4P-PDOT blocks the melatonin actions by selective Gq/11 G protein-coupled MT1 and MT2 receptors. Our data showed that melatonin treatment significantly reduced the accumulated lipid content in lipid droplets induced by the palmitate. It was through the melatonin receptors MT1 & MT2 and was confirmed by the treatment with melatonin receptor inhibitors luzindole and 4P-PDOT (Fig. 3). These inhibitors block the melatonin signaling by its receptors and hinder the positive effect of melatonin on lipid metabolism.

We further studied lipogenesis, adipogenesis, and lipolysis to identify the mechanism of lipid accumulation by palmitate and the impact of melatonin and its signaling. ACC1 gene is responsible for fatty acid synthesis, and SREBP1c and FASN regulate fatty acid synthesis and TAG esterification [36]. PPARγ acts as a transcription factor for various adipogenesis genes and is known as a master regulator [37].

A previous study suggests that the expression of lipogenesis/adipogenesis genes like SREBP1c, FASN, and PPARγ are increased by constant light exposure at night, whereas melatonin treatment decreases their expression [38]. Our results showed that melatonin treatment reduced the mRNA expressions of lipogenesis and adipogenesis genes like ACC1, FASN, SREBP1c (Fig. 4), and PPARγ (Fig. 5) in palmitate-induced conditions. The dysregulated lipogenesis induced by palmitate was restored to the control cell levels by melatonin, and the melatonin receptor inhibitors affect the positive effect of melatonin on dysregulated lipogenesis and adipogenesis. Therefore, we conclude that melatonin suppresses the de novo lipogenesis and adipogenesis by downregulating the SREBP1c, ACC1, FASN, and PPARγ via its MT1 and MT2 receptor signaling.

Melatonin rescues the cells from lipid accumulation induced by palmitate. Hence, we further explored melatonin and its receptor’s role in lipolysis as well. The ATGL and HSL are the rate-limiting enzymes of lipolysis. Previous research shows that the melatonin treatment to the bovine intramuscular preadipocytes reduced lipid droplet size and increased lipolysis genes like HSL, ATGL, and PLIN [39]. We report that the protein expression of ATGL (Fig. 6A) and HSL (Fig. 6B) was reduced by palmitate induction, and melatonin increased its expression (Fig. 6C) through its receptor signaling.

Obesity, in addition to dysregulated lipid metabolism and adipogenesis, elevates leptin secretion. Earlier studies reveal that excess leptin secretion leads to leptin resistance [40]. Another research shows that impaired melatonin signaling leads to leptin resistance [41]. It is reported that an increase in intracellular calcium ions can inhibit adipocyte differentiation or lipid accumulation. Cammisotto, P.G et al. [42] explained that increased calcium uptake and its release from endogenous stores inhibit insulin-stimulated leptin secretion in white adipocytes. In addition, hypoxia elevates leptin secretion in a calcium-dependent manner. In hypoxic conditions, a low calcium level increases leptin secretion, and a high calcium level inhibits excess leptin secretion [43]. Our result shows that the palmitate induction increases leptin secretion (Fig. 7) and reduces the intracellular calcium content (Fig. 8). Melatonin reduces elevated leptin secretion by increasing calcium release through the MT1 and MT2 receptors. Hence, the leptin secretion is regulated by intracellular calcium levels.

Activated melatonin receptors by melatonin inhibit the PKA/cAMP pathway. In the absence of PKA/cAMP signaling, the lipolysis is activated by other pathways like PI3K, PKC, and ERK [2]. Hence, the increase of lipolysis and decrease in lipogenesis, calcium release, and reduced leptin secretion by melatonin are activated by PI3K/AKT signaling. Inhibition of PKA/cAMP signaling by activating MT1 & MT2 receptors by melatonin leads to calcium release and activates PKC through PI3K/AKT signaling. A study shows that melatonin supplements to the palmitate-induced ovarian granulosa cells of polycystic ovary syndrome reduced insulin resistance by activating the PI3K/AKT pathway [44]. It is reported that chronic inflammation and ER stress caused by obesity disturb hepatic PI3K/AKT signaling (45).

A report suggests that the melatonin treatment in HFD-fed mice elevates the AKT protein expression [46]. Neuroprotection by melatonin was managed by its receptor signaling and activation of the PI3K/AKT pathway in primary astrocytes [47]. We observed that palmitate impairs PI3K (Fig. 9A) and AKT (Fig. 9B) mRNA expression, and melatonin treatment to the palmitate-induced cells increased the PI3K and AKT mRNA expression via its receptors. Hence, we report that melatonin exhibits its protective role in dysregulated lipid metabolism and leptin secretion by PI3K/AKT signaling through its receptors MT1 and MT2 in the palmitate-induced obese 3T3-L1 cell line.

## 5. Conclusion

In summary, melatonin, through its receptors, rescues the palmitate-induced cells from lipid accumulation in lipid droplets by reducing lipogenesis (SREBP1c, ACC1, FASN), adipogenesis (PPARγ) and increasing lipolysis, namely HSL and ATGL enzymes. In addition, melatonin reduces leptin secretion induced by palmitate by increasing intracellular calcium by activating MT1 and MT2 receptor signaling via the PI3K/AKT pathway.

## Abbreviations

4P-PDOT: 4-Phenyl-2-propionamidotetralin
ACC1: Acetyl-CoA carboxylase 1
ATGL: Adipose triglyceride lipase
ANOVA: Analysis of variance
BSA: Bovine Serum Albumin
cAMP: Cyclic Adenosine Monophosphate
CREB: cAMP Response Element-Binding Protein
cDNA: Complementary DNA
DMSO: Dimethyl Sulfoxide
DMEM: Dulbecco’s Modified Eagle Medium
ELISA: Enzyme-Linked Immunosorbent Assay
ERK: Extracellular Signal-Regulated Kinase
FASN: Fatty Acid Synthase
GPCR: G Protein-Coupled Receptor
HSL: Hormone-Sensitive Lipase
IgG: Immunoglobulin G
IP_3_: Inositol trisphosphate
kcal/day: Kilocalories per day
MT1: Melatonin Receptor 1
MT2: Melatonin Receptor 2
MTT: 3-(4,5-Dimethylthiazol-2-yl)-2,5-Diphenyltetrazolium Bromide
PPARγ: Peroxisome Proliferator-Activated Receptor Gamma
PGC1a: Peroxisome Proliferator-Activated Receptor Gamma Coactivator 1-alpha
PBS: Phosphate Buffered Saline
Pi3k: Phosphoinositide 3-kinase
PLA2: Phospholipase A2
PLC: Phospholipase C
PKA: Protein Kinase A
PKC: Protein Kinase C
RIPA: Radioimmunoprecipitation Assay
SREBP1c: Sterol Regulatory Element-Binding Protein 1c
TAG: Triacylglycerol

## Acknowledgment

We thank the Department of Science and Technology, Government of India, for the fellowship under Women Scientist Scheme-A, UGC-BSR Fellowship, UGC, Government of India, and acknowledge the instrument facilities from the Department of Biochemistry (under DST-FIST program), Centre for Excellence from the School of Life Sciences and DST-PURSE program of Bharathidasan University, Tiruchirappalli, India.

